# Using Machine Learning to Parse Breast Pathology Reports

**DOI:** 10.1101/079913

**Authors:** Adam Yala, Regina Barzilay, Laura Salama, Molly Griffin, Grace Sollender, Aditya Bardia, Constance Lehman, Julliette M. Buckley, Suzanne B. Coopey, Fernanda Polubriaginof, Judy E. Garber, Barbara L. Smith, Michele A. Gadd, Michelle C. Specht, Thomas M. Gudewicz, Anthony Guidi, Alphonse Taghian, Kevin S. Hughes

## Abstract

1

**Purpose:** Extracting information from Electronic Medical Record is a time-consuming and expensive process when done manually. Rule-based and machine learning techniques are two approaches to solving this problem. In this study, we trained a machine learning model on pathology reports to extract pertinent tumor characteristics, which enabled us to create a large database of attribute searchable pathology reports. This database can be used to identify cohorts of patients with characteristics of interest.

**Methods:** We collected a total of 91,505 breast pathology reports from three Partners hospitals: Massachusetts General Hospital (MGH), Brigham and Womens Hospital (BWH), and Newton-Wellesley Hospital (NWH), covering the period from 1978 to 2016. We trained our system with annotations from two datasets, consisting of 6,295 and 10,841 manually annotated reports. The system extracts 20 separate categories of information, including atypia types and various tumor characteristics such as receptors. We also report a learning curve analysis to show how much annotation our model needs to perform reasonably.

**Results:** The model accuracy was tested on 500 reports that did not overlap with the training set. The model achieved accuracy of 90% for correctly parsing all carcinoma and atypia categories for a given patient. The average accuracy for individual categories was 97%. Using this classifier, we created a database of 91,505 parsed pathology reports.

**Conclusions:** Our learning curve analysis shows that the model can achieve reasonable results even when trained on a few annotations. We developed a user-friendly interface to the database that allows physicians to easily identify patients with target characteristics and export the matching cohort. This model has the potential to reduce the effort required for analyzing large amounts of data from medical records, and to minimize the cost and time required to glean scientific insight from this data.

## 2 Introduction

Pathology reports, individually and in the aggregate, contain immense quantities of information critical to cancer research. Unfortunately, the majority of this information is present as free text, locking the data away from all but the most tenacious researcher. It is extremely difficult to parse out useful data from the free text without significant work by human beings. Therefore, when studies are undertaken, they involve a step of manually extracting relevant information from free-text reports and coding it into structured data.

This makes large-scale studies prohibitively expensive as they require well-educated and trained individuals, often at the MD level, to participate in data preparation. Automating this process would not only reduce costs, but would also enable the research community to apply big data analysis methods to clinical research and population based patient care.

Natural language processing (NLP) is a computerized approach for extracting the meaning within free-text, translating prose into a structured database that contains information of interest. There are basically two approaches to NLP, a rules based approach, where words and phrases within free-text are mapped to categories, and a machine learning approach, where a training set of annotated reports are provided to a computer which learns how to identify desired categories.

Rule-based NLP techniques have been applied to breast pathology reports with varying results. For instance, Buckley et al [1] designed a set of rules for mapping pathology reports to values of extracted fields. The manual efforts involved in designing such rules turned out to be labor intensive and not easily extended to other organs or disease processes. For instance, [1] showed that invasive ductal carcinoma (IDC) had been stated in 124 different ways in their corpus of pathology reports. Similar findings have been reported by others who employed rule-based approaches for the task [1, 11, 5, 2, 8, 6].

An alternative approach to text processing is to employ machine learning techniques. Instead of using manually specified rules, these approaches learn extraction patterns from pairs of pathology reports with associated values. Typically, the accuracy of these algorithms depends on the size of available annotations. However, in the case of pathology reports, such annotations are commonly readily available from previous clinical studies, and can be re-purposed. A number of researchers employed these techniques for parsing breast pathology reports [4, 3, 7, 12]. Unfortunately, the reported results have been unsatisfactory. For instance, a recent paper [12] achieved accuracy of 12.6%.

In this study, we apply a modern machine learning technique with a large annotated training set to automatically parse free-text pathology reports into structured data. In contrast to prior approaches, our model achieves excellent performance with 90% per report accuracy and 97% per category accuracy, and requires only a few development days.

### 3 Methods

#### Pathology Reports

With the approval of the Partners Institutional Review Board (IRB), we collected a total of 102,907 breast pathology reports from three Partners hospitals: Massachusetts General Hospital (MGH), Brigham and Women's Hospital (BWH), and Newton-Wellesley Hospital (NWH), covering the period from 1978 to 2016. The pathology reports in the three Partners institutions do not follow the same structure and exhibit marked differences in verbalizations.

Using the College of American Pathologists' (CAP) synoptic reporting system as a starting point, we identified 20 categories of information that were known to be useful in categorizing breast disease and breast cancer (shown in Table 2).

The work-flow for processing these reports was as follows: First, all reports were automatically anonymized, stripping dates, medical record numbers, and patient names from the free-text of each pathology report as shown in the left-hand side of Figure 2. Second, 11,402 non-breast pathology reports were identified and excluded, leaving 91,505 reports breast pathology reports. Third, reports of bilateral breast cases were split into two reports and parsed into two records in the database (one right and one left). This resulted in a database with 108,114 rows.

Machine learning requires an annotated dataset where free-text reports have been annotated for the correct value of each category. We aggregated the annotations in our dataset from two sources (both of which covered subsets of the records in the above mentioned database: 1) The dataset used by Buckley et al [1] which included annotations for a nonrandom sample of 6,295 reports over 3,313 patients annotated for various types of carcinoma and atypia. 2) The Breast Cancer Database (BCD) Partners IRB: 1999P009256 which includes annotations for 10,841 reports over 5,268 patients about typical tumor characteristics (categories) such as estrogen receptor (ER), progesterone receptor (PR) and HER2 status as well as carcinomas and atypias.

There was some overlap in the categories covered in the two sources and different categories had widely varying amounts of annotation. For example, the Isolated Tumor Cells (ITC) category has 1,143 annotations from only the BCD, and the Ductal Carcinoma In Situ (DCIS) category had 16,550 annotations, which come from a combination of both sources. Table 2 shows the full list of categories, how much annotation was available per category, and their respective sources.

For each category, we split out our annotated dataset into a training set and a test set. The training set was the set of reports and respective annotations that we gave to our machine learning model to learn from. The test set was the held-out set of reports and annotations that we used to evaluate our model's predictions. We note that the train/test sets for all 20 categories were independent, and there was a separate training set and test set for each category. In general, we used 500 annotations for our test set, with the exception of the Extra Capsular Extension (ECE), and ITC categories, for which we used 203 and 171 held out annotations. This was done because we had relatively little annotation for those categories (1,354 and 1,153 respectively), and so we used 15% of available annotations as testing sets instead of 500 as in other data-rich categories.

#### Machine Learning Method

The training set was then used to train our classifier using boosting classification, where a strong non-linear classification was achieved by combining weak learners, such as decision stumps [10]. We represented each pathology report using standard n-gram representation, where the text was a vector capturing words and phrases that appeared in the document. During training, the model learned the weights of each phrase. We trained the classifier separately for each one of 20 categories.

#### Evaluation Methodology

For evaluation, we compared our models predictions with the annotations of our held-out test set.

We calculated Accuracy, Precision, Recall (Sensitivity), and F1 score for the possible values of each category (e.g. Positive, Negative or Not available). Accuracy was defined as the portion of times the models prediction agreed with the annotations. Precision was defined as the True Positive rate (what portion of the positives we predicted were correct). Recall was defined as the proportion of all true positives our model captured. F1-score was the harmonic mean of Precision and Recall. F1-score ranged from 0 to 1, where 1 was perfect performance. To achieve a high F1-score our model had to both predict positives very accurately (Precision) and miss very few positives (Recall).

In addition to per-category analysis, we reported “all-or-nothing” report-level accuracy for the major categories of carcinoma and atypia, all of which can either be Present or Absent. For our model to be correct in “all-or-nothing” evaluation, it had to predict all categories of carcinoma and atypia correctly.

#### Comparison with Rule-Based Approach

To compare with the previously reported rules based approach [1], we manually examined reports where our models disagreed. We randomly selected 15 reports for each of the 9 shared categories, resulting in 135 reports overall. The list of shared categories can be seen in Table 2.

#### Learning curve analysis

We also performed a Learning Curve analysis, where we plotted the performance of the system, measured in aggregate F-score over the cancer and atypia categories, as we varied the amount of annotation from 10 examples to 5,500 examples. This type of analysis lets us analyze how much annotation our model needed to start achieving reasonable performance.

#### Creating Database

After training our model, we applied it to the full set of 108,114 free-text preprocessed reports to predict the values of all 20 categories. To make this accessible, we have made this resource available to other researchers through an internal web-interface, as shown in Figure 1, which was accessible with IRB approval.

**Figure 1:**
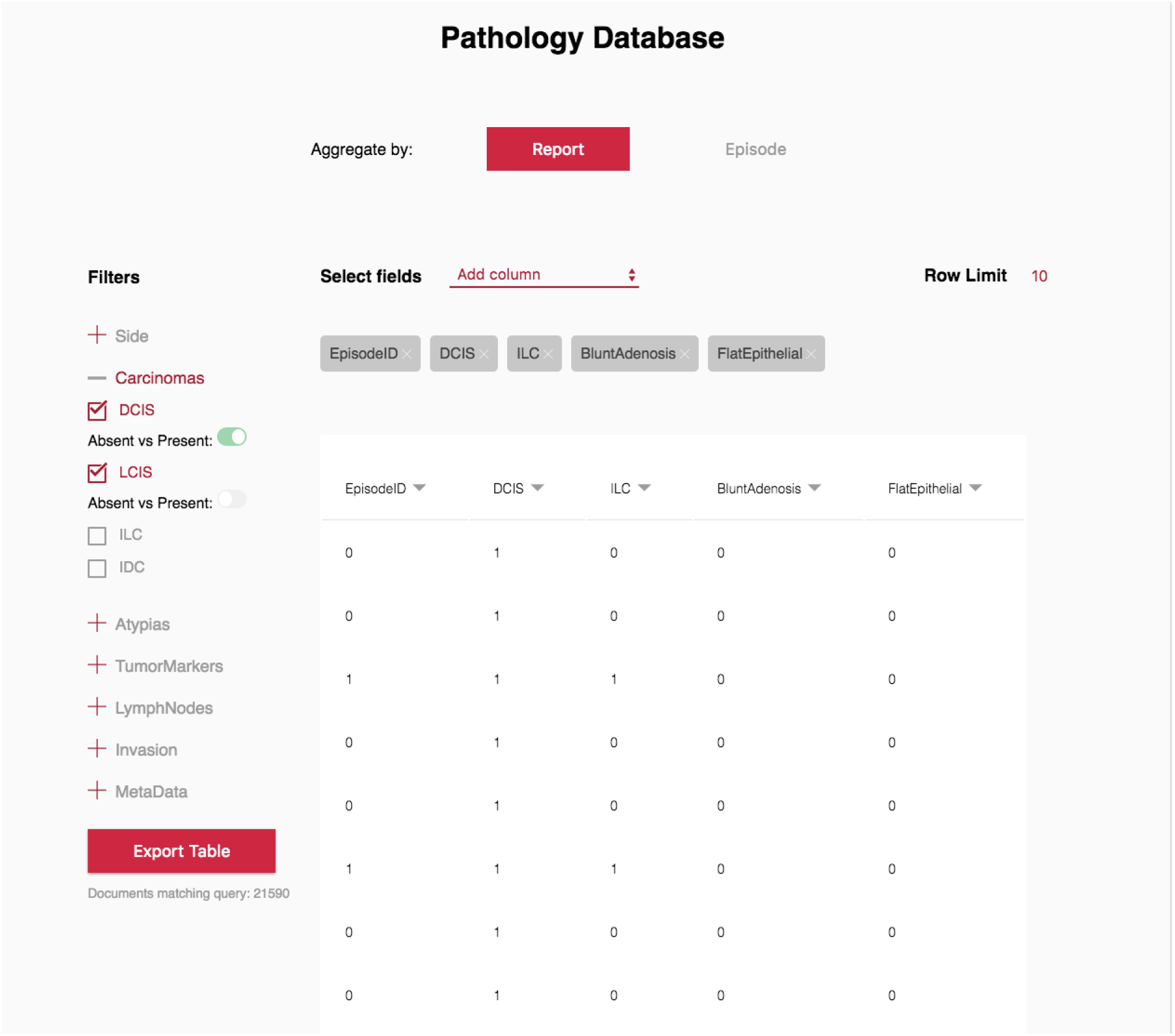
A screenshot of the searchable database we built on the results of our system. Users can query for any combination of diagnoses and export the resulting database

In addition to report-based retrieval, we also enabled the users to perform *episode-based* search. In managing breast patients, it was not unusual to have multiple pathology reports relating to a single episode of care as we previously described [1]. An episode of care included all pathology reports for a single side within a six-month period. For example, a patient might have a core biopsy showing atypical ductal hyperplasia (ADH), followed by an excision showing ductal carcinoma in situ (DCIS), followed by a lumpectomy showing invasive ductal carcinoma (IDC). This information would be consolidated into a single episode showing that the patient at that time had ADH, DCIS and IDC.

### 4 Results

An example of system input and output is shown in Figure 2. We evaluated our performance by comparing our model's predictions on a held out set of 500 pathology reports for each category against their corresponding MD annotations.

**Figure 2:**
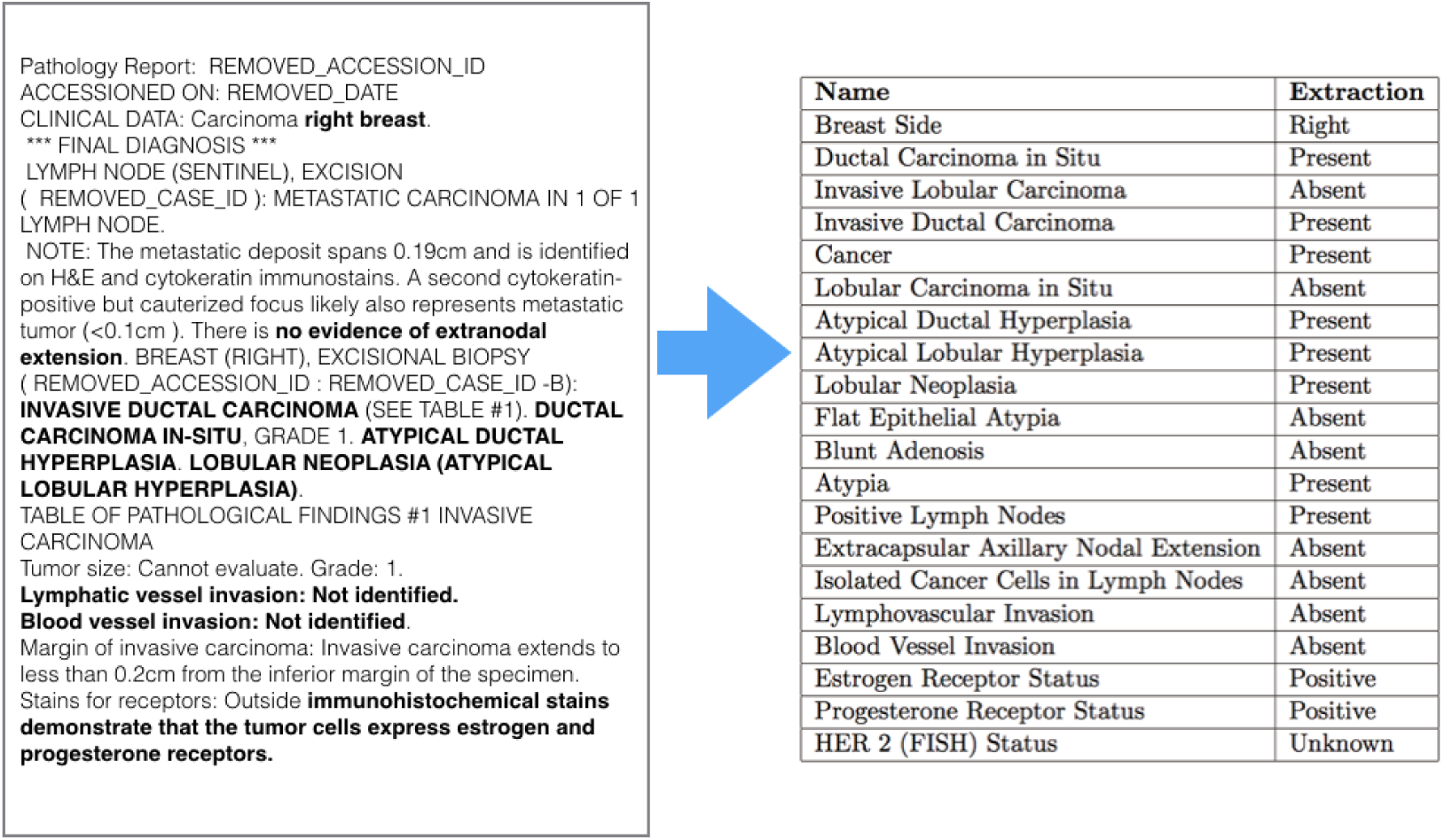
An example of an anonymized pathology report and the extractions for that report. We bold the text in the report relevant to our extractions for clarity.

#### Data statistics

The average length of a pathology report after preprocessing was 354 words, with the shortest being 17 words and the longest being 2,428 words. Statistics on pathology report lengths broken out by Institution can be seen in Table 1.

**Table 1:**
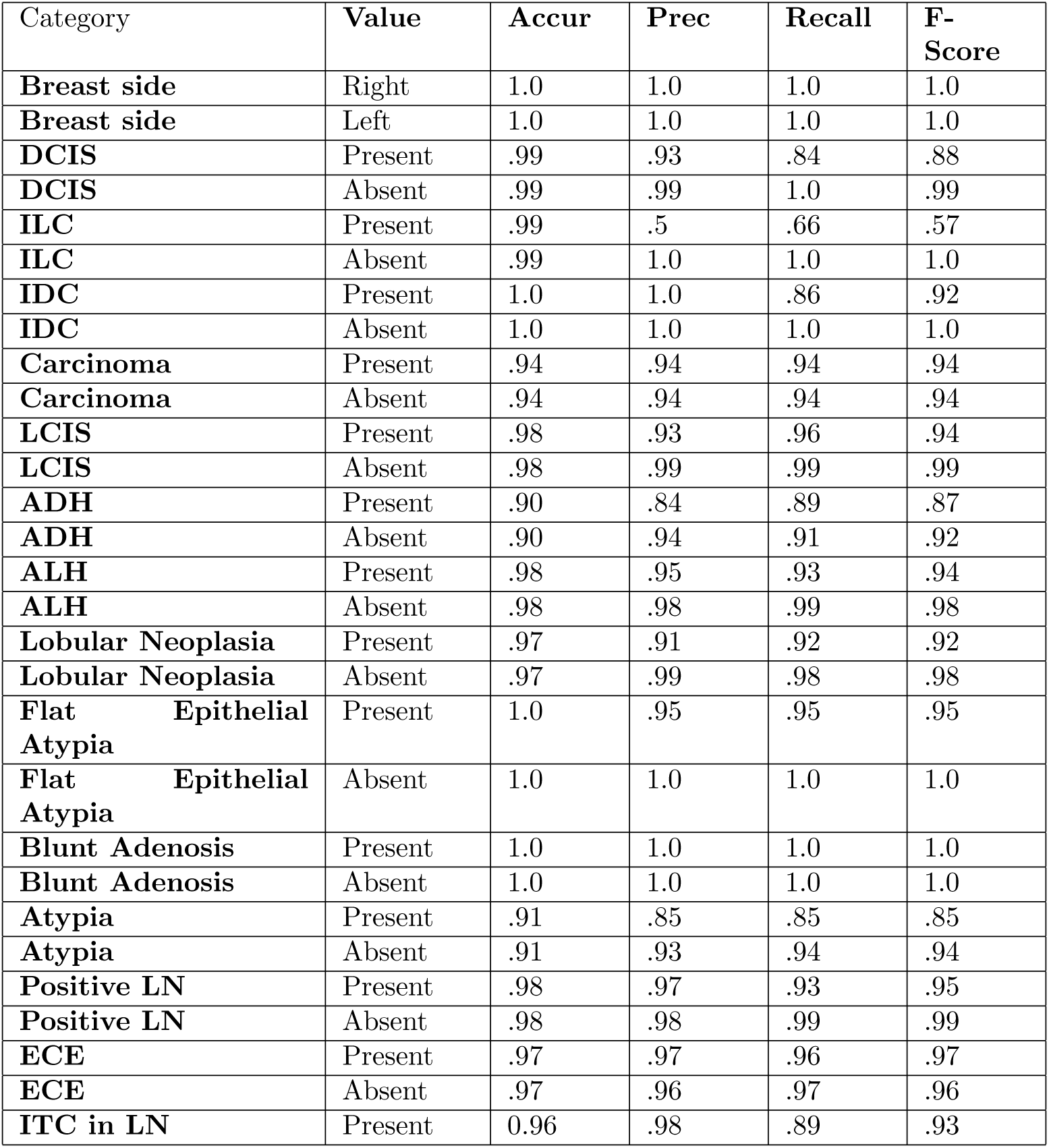
Statistics on pathology reports broken out by institution, after preprocessing

**Table 2:**
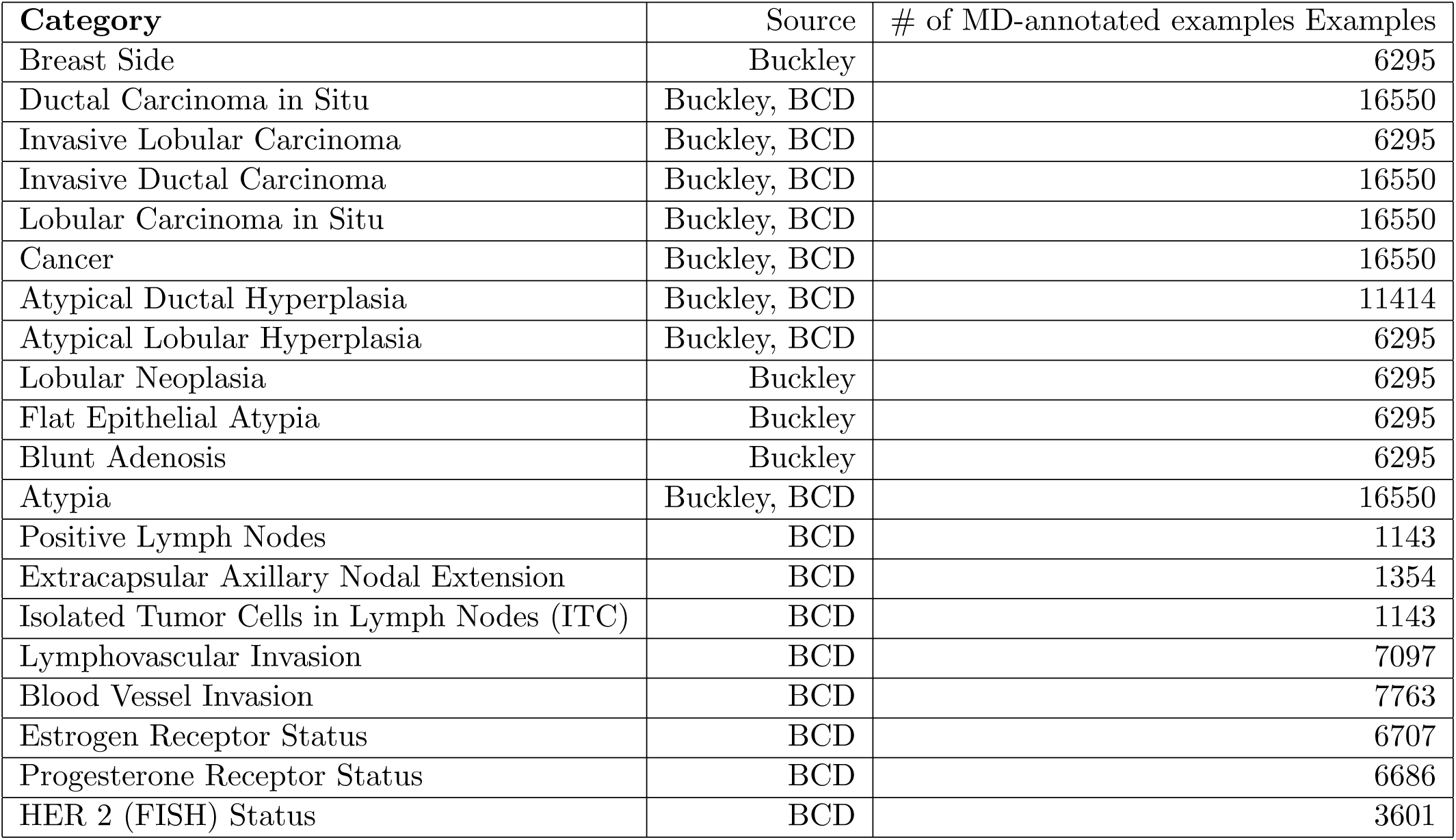
Source and amount of labeled examples and for each category. Buckley refers to [1], and BCD refers to the Breast Cancer Database (Partners IRB: 1999P009256).

#### Per category performance

Table 3 displays extraction accuracy, precision, recall and F-score for all possible values of all 20 categories. Table 4 shows the aggregate accuracy and F-score for each category. The latter table shows that 19 out of 20 categories have F-scores greater than .90. The remaining category has an F-score of 0.88. All categories had an accuracy over 90%, and 16 categories had an accuracy over 95% on our test set. Our model had an average category F-score of 0.96 and an average category accuracy of 97%.

**Table 3:**
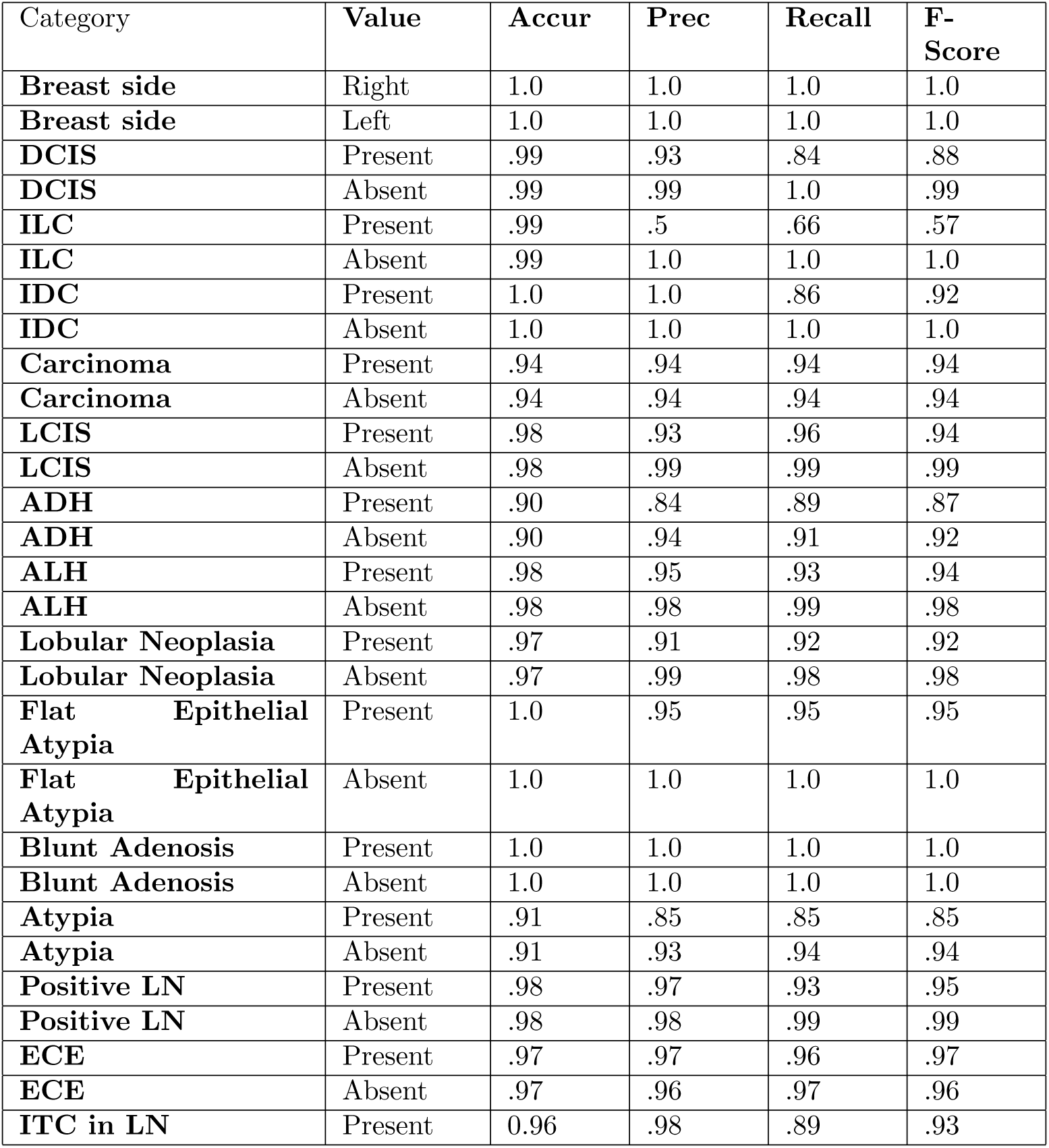
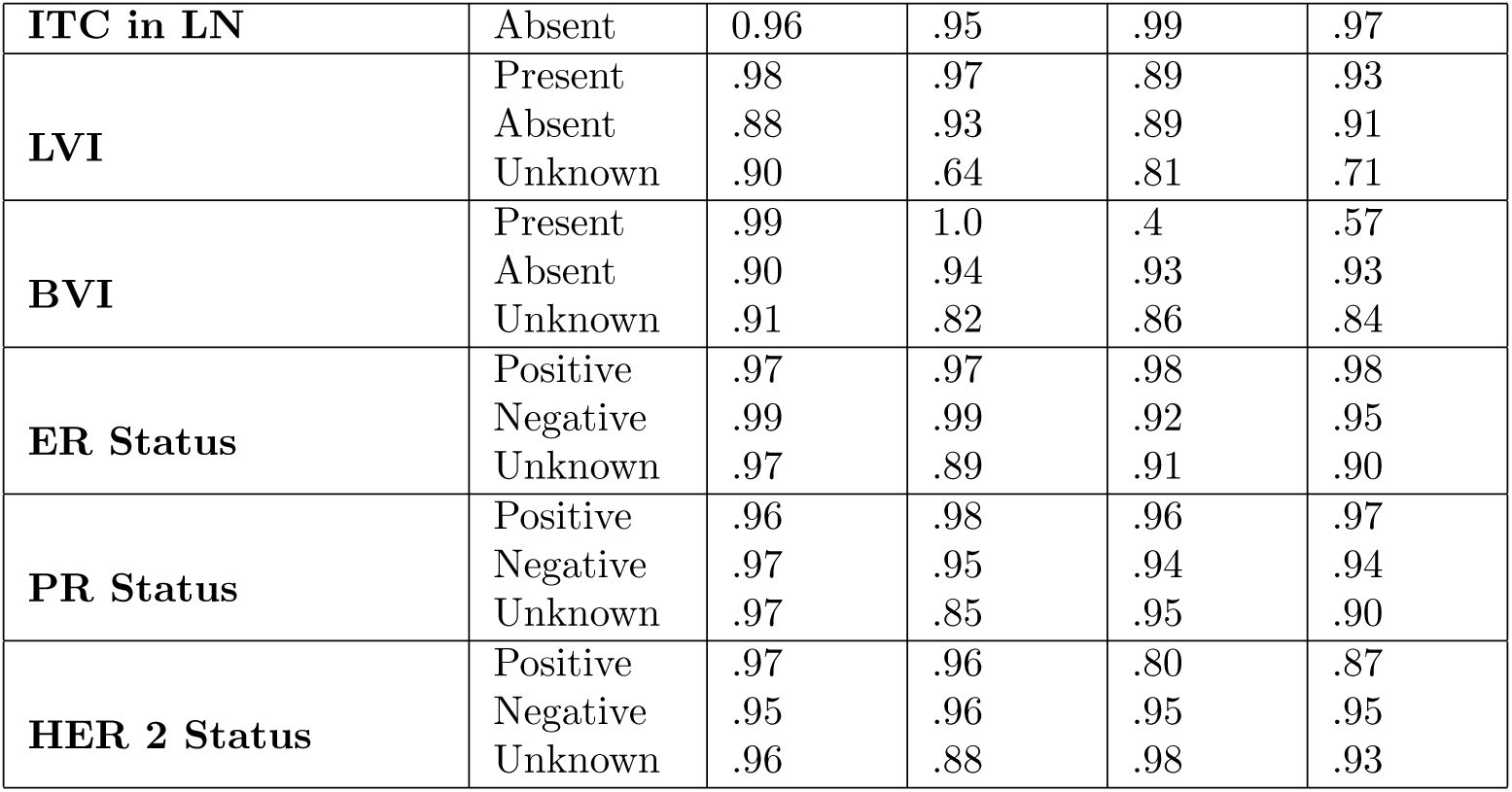
This tables shows Accuracy, Precision, Recall and F-Score for each value of each category. All these results come from comparing the predictions of our model to MD annotations on a heldout test set of 500 reports. We note that for ECE, and ITC, we present evaluate our on 203 and 171 held out reports respectively, instead of 500, because we have relatively few annotations for those categories.

**Table 4:** This table shows F-score and accuracy for each category, the average across all categories and “All-or-nothing” report-level accuracy.

| Category | Accuracy | F-score |
| --- | --- | --- |
| **Breast side** | 1.0 | 1.0 |
| **DCIS** | .99 | .99 |
| **ILC** | .99 | .99 |
| **IDC** | 1.0 | 1.0 |
| **Carcinoma** | .94 | .94 |
| **LCIS** | 1.0 | .98 |
| **ADH** | .90 | .90 |
| **ALH** | .98 | .98 |
| **Lobular Neoplasia** | .97 | .97 |
| **Flat Epithelial Atypia** | 1.0 | 1.0 |
| **Blunt Adenosis** | 1.0 | 1.0 |
| **Atypia** | .91 | .91 |
| **Positive LN** | .98 | .98 |
| **ECE** | .97 | .97 |
| **ITC in LN** | .96 | .96 |
| **LVI** | .92 | .88 |
| **BVI** | .93 | .90 |
| **ER Status** | .97 | .96 |
| **PR Status** | .97 | .95 |
| **HER 2 Status** | .96 | .94 |
| **Report-Level** | .90 | N/A |
| **Average** | .97 | .96 |

We note that the model predicted *carcinomas* with an F-score 0.94 and an accuracy of 94%. Our model predicted the *atypias* with an F-score of 0.91 and an accuracy of 91%.

#### Per report Performance

We also computed the “all-or-nothing” report-level accuracy where the system must correctly extract all the information about atypias and carcinomas from a given report to be correct. In this evaluation, the system achieved 90%. Since this evaluation required correct prediction across multiple categories at once, the performance was lower than per-category accuracy. However, it was still sufficiently high to be used in practical setting.

#### Comparison with rule-based approach

To compare to the rule-based model presented in [1], we selected 135 random reports where the rule-based model and our model disagreed, 15 from each of the 9 shared categories. We manually compared the 135 disagreements and observed that our model was correct 52% of the time, showing our model performed as well as the previous work while requiring much less manual effort.

#### Learning curve analysis

Our next question concerned the amount of annotations required for achieving satisfactory performance. This was a pertinent question because of the cost associated with obtaining a large corpus of manual annotations. Figure 3 shows the average F1 score as we varied the training set size from 10 annotations per category to 5,500. With only 400 annotations, the model achieved an average category F-score of 0.90. This result shows the promise of using this approach in annotation-lean scenarios.

**Figure 3:**
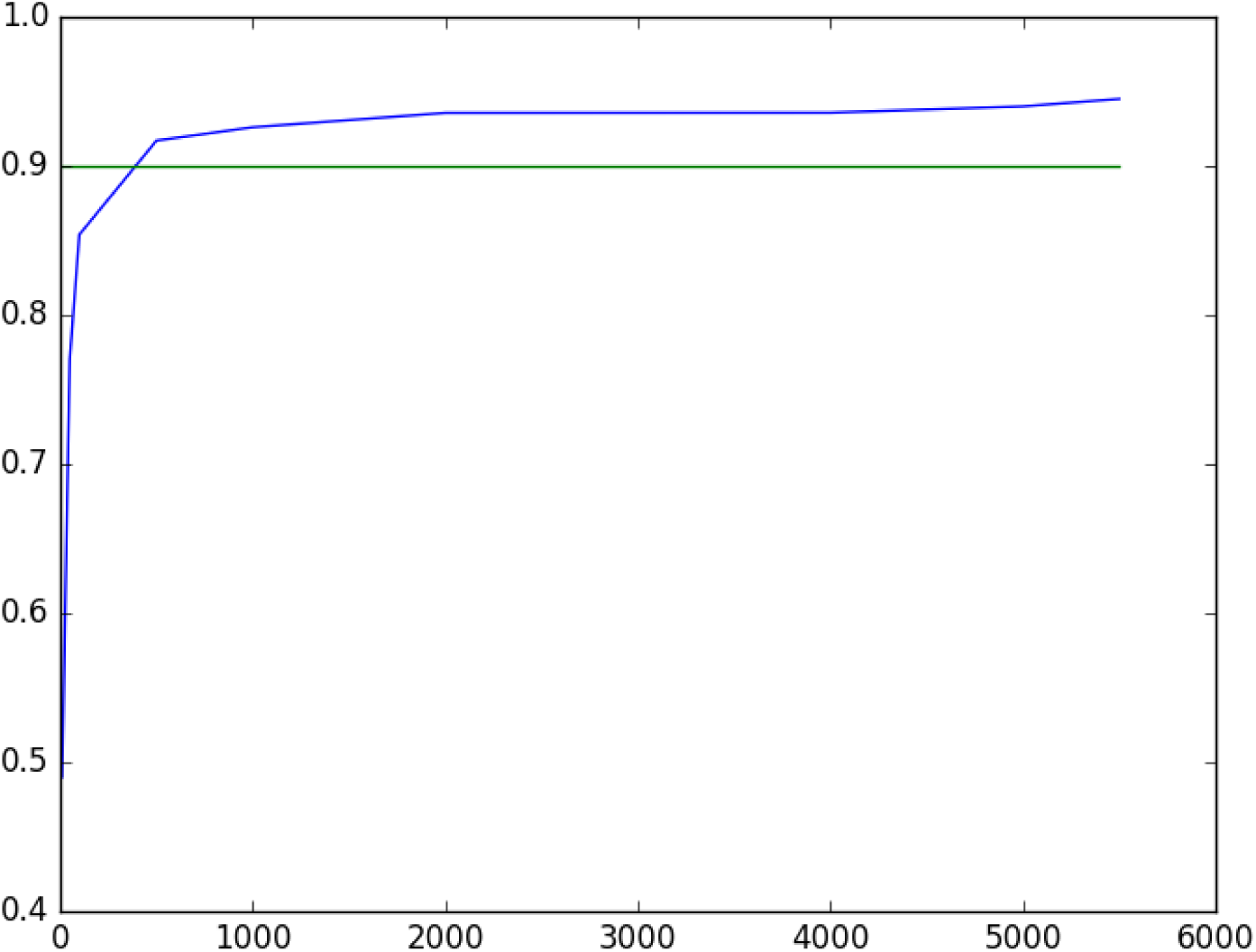
The average F1-score of our model plotted against the amount of supervision, ranging from 10 to 5,500 annotations per type of information

#### Output

Using 91,505 pathology reports for 49,717 patients, we created a database having 108,114 rows (records) which could be consolidated to 73,542 episodes of care. An example of system input and output is shown in Figure 2.

### 5 Discussion

Our results demonstrate that machine-learning system can accurately parse breast pathology reports rivaling the performance of a carefully engineered rule-based system [1]. Our approach saves significant manual effort involved in manual engineering employed by the rule-based approaches. As stated by Buckley et al [1]:

> “We have been struck by the inherent complexity of using NLP in medical care. The time and effort required to use NLP for a single, well-defined problem should give pause to the idea that having data in any electronic format, even free text, will help us improve medical care.”

At the same time, we show significant improvement over previous machine learning approaches. Of these approaches, Wieneke et al [12] is the closest to our work. Their extraction scheme has a large overlap with our categories, including features such as Ductal Carcinoma in Situ (DCIS), Lobular Carcinoma in Situ (LICS), and Atypical Lobular Hyperplasia (ALH), and they rely on a medium-sized training set of 3,234. However, their model achieves low performance for most categories, with an F-Score of 0 for DCIS, LCIS, and ALH. Other researchers report improved performance on different annotation schemes e.g. 0.65 [3], 0.82 [7] and 0.85 [9].

In this paper, we demonstrate that machine learning techniques can accurately parse pathology reports reaching an F-score of 0.96, without daunting manual effort. We hypothesize that the success of our method can be attributed to the learning algorithm and the quality of the annotated data. The boosting learners are known to achieve top performance on the classification tasks, and can robustly handle the noise in the training data. The annotations used for our algorithm contain a large representative sample of diverse pathology reports from three different hospitals, and are carefully annotated by trained professionals.

To make our system results usable in a clinical setting, we have compiled parsed pathology reports into a database with a user-friendly interface, as shown in Figure 1. This interface can be used by researchers with appropriate IRB approval to identify cohorts of patients determined by specific combinations of categories. These cohorts can be identified as single records or as episodes of care, which is defined as a consolidated record that contains all pathology categories found within a six month period on the same side (left or right) [1]. For example, a researcher looking for pure DCIS cases would want to search by episode of care, not by specific report. Once a cohort is identified, all pathology reports in the database for each individual in the cohort can be retrieved. This provides a quick way to analyze prior diseases for each patient (which might exclude the patient from a study) or subsequent diagnoses (potentially an endpoint of a study). For example, a researcher can see the risk of subsequent cancer in patients with high risk lesions while excluding patients with prior cancers or can see the rate of in-breast or contralateral disease, giving a first pass at the long term clinical course. Obviously patients change health care systems regularly, so while this first pass will not give comprehensive follow up information, it will provide a valuable start in gathering the data for the research endeavor.

The cohort and all pathology reports over time are made available to the researcher, who must then review the records for accuracy, since we did not achieve an F-score of 1. As each record is reviewed and edited, the corrected records are returned and added to the training set, which is used to further enhance the algorithm. Over time, we expect that as multiple records are corrected and used for retraining of the algorithm, we will asymptotically approach an F-score of 1.

We are also exploring the possibility of making this system publicly available, allowing the uploading of de-identified pathology reports from other institutions to be parsed by our algorithm. While we expect F-scores to be lower as reports from different institutions may have different formats or linguistic creations, we also expect that this will improve over time. It should also be noted that the machine learning approach described here can be applied to any language with annotated reports, as the algorithm looks for patterns as they relate to annotations, not at actual words or phrases. We are currently exploring this approach for non-English reports.

We recognize that our system has a number of limitations. Our algorithm is not 100% accurate and thus requires post-processing before use for clinical care or research. Compared to the absence of this data for large-scale studies or clinical care, we feel this is acceptable, but at the same we are developing workflows that we anticipate will markedly increase our accuracy. We also recognize that our algorithm might not be immediately applicable to other institutions. We are addressing this by possibly opening our system for general use, where annotated reports and corrections will strengthen our system for all institutions. As we parse annotated data from more and more institutions, we expect our algorithm will learn to be more generalizable, as there is some basic similarity of breast pathology reports and a limit to how creative pathologists can be in describing the same disease scenario.

### 6 Conclusion

We have developed a robust system for NLP of breast pathology reports that we anticipate will become more accurate and useful over time. We are hopeful that this system will be useful for research and clinical care at our institution and potentially at multiple institutions. We are also hopeful that this approach can be extrapolated to other free-text medical reports, hopefully salvaging at least some of the massive amounts of information locked in the free-text of the electronic health records.

